# P2X-GCaMPs as versatile tools for imaging extracellular ATP signaling

**DOI:** 10.1101/2020.04.23.057513

**Authors:** Matthias Ollivier, Juline Beudez, Nathalie Linck, Thomas Grutter, Vincent Compan, Francois Rassendren

## Abstract

Adenosine 5’ triphosphate (ATP) is an extracellular signaling molecule involved in numerous physiological and pathological processes. Yet, *in situ* characterization of the spatiotemporal dynamic of extracellular ATP is still challenging due to the lack of sensor with appropriate specificity, sensitivity and kinetics. Here we report the development of biosensors based on the fusion of cation permeable ATP receptors (P2X) to genetically encoded calcium sensors (GECI). By combining the features of P2X receptors with the high signal to noise ratio of GECIs, we generated ultrasensitive green and red fluorescent sniffers that detect nanomolar ATP concentrations *in situ* and also enable the tracking of P2X receptor activity. We provide the proof of concept that these sensors can dynamically track ATP release evoked by neuronal depolarization or by extracellular hypotonicity. Targeting these P2X-based biosensors to diverse cell types should advance our knowledge of extracellular ATP dynamics *in vivo*.

## Introduction

Adenosine 5’ triphosphate (ATP) is an ubiquitous molecule that provides cellular energy to all living organisms. In most species, except in insects and nematodes (Hou and Cao, 2016), ATP is also an extracellular messenger that allows cell-to-cell signaling through activation of specific purinergic plasma membrane receptors (Burnstock, 2018). ATP signaling is involved in the regulation of plethora of physiological functions and increasing number of evidence supports that purinergic signaling components are remodeled in pathological conditions, thereby directly contributing to diseases manifestation (Burnstock, 2006).

There are two main families of membrane receptor for ATP, the P2X and P2Y receptors. P2X receptors are ATP-gated channels formed by the association of three subunits (Kawate et al., 2009; Khakh and Alan North, 2006). The seven P2X subunits can form homo- or heteromeric channels, the later with different stoichiometries (Compan et al., 2012). Despite their identification more than twenty years ago, pharmacology of P2X receptors still remains limited. Indeed, because of their widespread expression, of our partial understanding of their subunit composition and the relative paucity of pharmacological tools, deciphering the physiology of P2X receptors is still challenging. Extracellular ATP also activates a subset of G protein coupled P2Y receptors (von Kügelgen and Harden, 2011), mainly P2Y1, P2Y2 and P2Y11, although these receptors display higher affinity for other endogenous nucleotide such as ADP or UTP (von Kügelgen, 2019). Yet, activation of P2Y receptors by ATP seems to have important physiological functions, particularly in the central nervous system (A. Weisman et al., 2012). Pharmacology of P2Y receptors is considerably more developed than that of P2X receptors, particularly for P2Y receptors that are sensitive to ATP (Jacobson and Müller, 2016). However, because these proteins are coupled to intracellular signaling pathways, studying the dynamic of their activation in integrated preparation mostly relies on indirect readthrough.

Studying extracellular ATP signaling in multicellular preparations and *in vivo* is complex. First, in vertebrate virtually all cells can release ATP through various mechanisms such as classical vesicular release, lysosomal exocytosis, transmembrane channels or cell lysis (Lohman et al., 2012; Praetorius and Leipziger, 2009; Rassendren and Audinat, 2016). Signals triggering ATP release remain poorly characterized, although there is clear evidence that cells constitutively release ATP (Lazarowski et al., 2011; Sivaramakrishnan et al., 2012) and that evoked release can be triggered in defined physiological and pathological contexts (Lazarowski, 2008). A second level of complexity comes from the short half-life of extracellular ATP. Due to the ubiquitous expression of membrane-bound and soluble ectonucleotidases, extracellular ATP is within seconds to minutes degraded in ADP and ultimately adenosine, which both also act as signaling molecules (Deaglio and Robson, 2011; Kukulski et al., 2011).

Various methods have been developed to measure extracellular ATP (Wu and Li, 2020). The most common is the luciferase-luciferin assay which allows bioluminescent quantification of ATP in solution. A genetically encoded version of this assay was developed (Pellegatti et al., 2005), however because of its low quantic yield, this approach has limited applications and lacks spatio-temporal resolution. To assess ATP release *in situ*, electrochemical-based microelectrodes have been engineered. While these approaches provide a direct monitoring of extracellular ATP *ex vivo*, their spatial definition remains limited (Llaudet et al., 2005). Several genetically encoded extracellular ATP sensors have been engineered (Conley et al., 2017; Lobas et al., 2019; Pellegatti et al., 2005; Richler et al., 2008). They present several improvements to chemical-based assays by allowing precise cellular and plasma membrane targeting as well as bioluminescent or fluorescent output. However, these sensors have several limitations that restrict their use. First, these sensors display poor apparent affinity for ATP, often above 100 μM and therefore are not sensitive enough to detect ATP concentration below 10 μM. Second, their kinetics are slow with τ-on and τ-off above 5 seconds at physiological ATP concentrations. These limitations considerably restrict their use *ex*- and *in-vivo* where extracellular ATP rarely raises above ten of micromolar and is rapidly degraded by ectonucleotidases.

To overcome these limitations, we generated versatile P2X-GCaMP6s fusion proteins that allow reporting accurately, specifically and dynamically their activity. In addition, engineering gain-of-affinity mutations in P2X2-GCaMP6s led to generate ATP sensors with nanomolar apparent affinity for ATP and fast kinetics.

## Results

P2X2-GCaMP6s fusion (stated below as PG6) was engineered by fusion of GCaMP6s, a genetically encoded calcium indicator (GECI) with improved kinetics, dynamic range and affinity for calcium (Chen et al., 2013) to the C-terminal tail of rat P2X2. We expected that upon receptor activation, calcium influx through P2X2 pore will specifically trigger GCaMP6s fluorescence as previously described using a FRET-based GECI (Richler et al., 2008) (Fig. 1A). Functional properties of P2X2 and PG6 were compared by whole cell recording in transfected HEK cells. We observed no difference in the apparent affinity for ATP of the two receptors or in their kinetics of activation and inactivation, although these later parameters were not investigated thoroughly (Fig. 1B, C). Upon 10 μM ATP application a strong increase of fluorescence near the plasma membrane was observed in HEK cell transfected with PG6 (Fig. 1D), while basal fluorescence was very low and did not change upon application of vehicle. We next used a plate reader assay combined with fast automated injection system to further characterize ATP-evoked fluorescence in PG6 transfected HEK cells. ATP dose-response experiment revealed an EC50 of 2.4 ± 0.3 μM (N = 4) and nH of 1.7 ± 0.2 with a dF/F of 1.3 ± 0.3 at 10 μM ATP (Fig. 1E and 1J). In our conditions of drug application and signal acquisition, analysis of activation time constants shows concentration dependent relationship approaching 1 second at saturating ATP concentration (Fig. 1F), although these values are certainly underestimated due to the frequency acquisition of the plate reader (1 Hz). Calcium influx through the P2X2 channel was responsible for GCaMP6s fluorescence, since removal of extracellular calcium completely abolished ATP-evoked signal (Fig. 1G). The contribution to fluorescence signal by calcium released from intracellular stores following activation of endogenous P2Y receptors by ATP was evaluated in HEK cells transfected with P2X2-K69A-GCaMP6s (stated bellow as PKG6), a biosensor in which the lysine 69 that contributes to the ATP-binding site was mutated to alanine (Jiang et al., 2000). In these cells, application of increasing ATP concentrations only evoked a minimal calcium signal (Fig. 1H and 1J). By comparison, in HEK cells expressing a cytosolic GCaMP6s, ATP-evoked endogenous P2Y activation resulted in a strong fluorescence signal with an EC50 of 10.6 ± 4 μM, nH = 1.5 (N = 4) (Fig. 1I and 1J), supporting that fluorescence signal evoked by PG6 activation is minimally contaminated by calcium release from internal stores.

**Figure 1:**
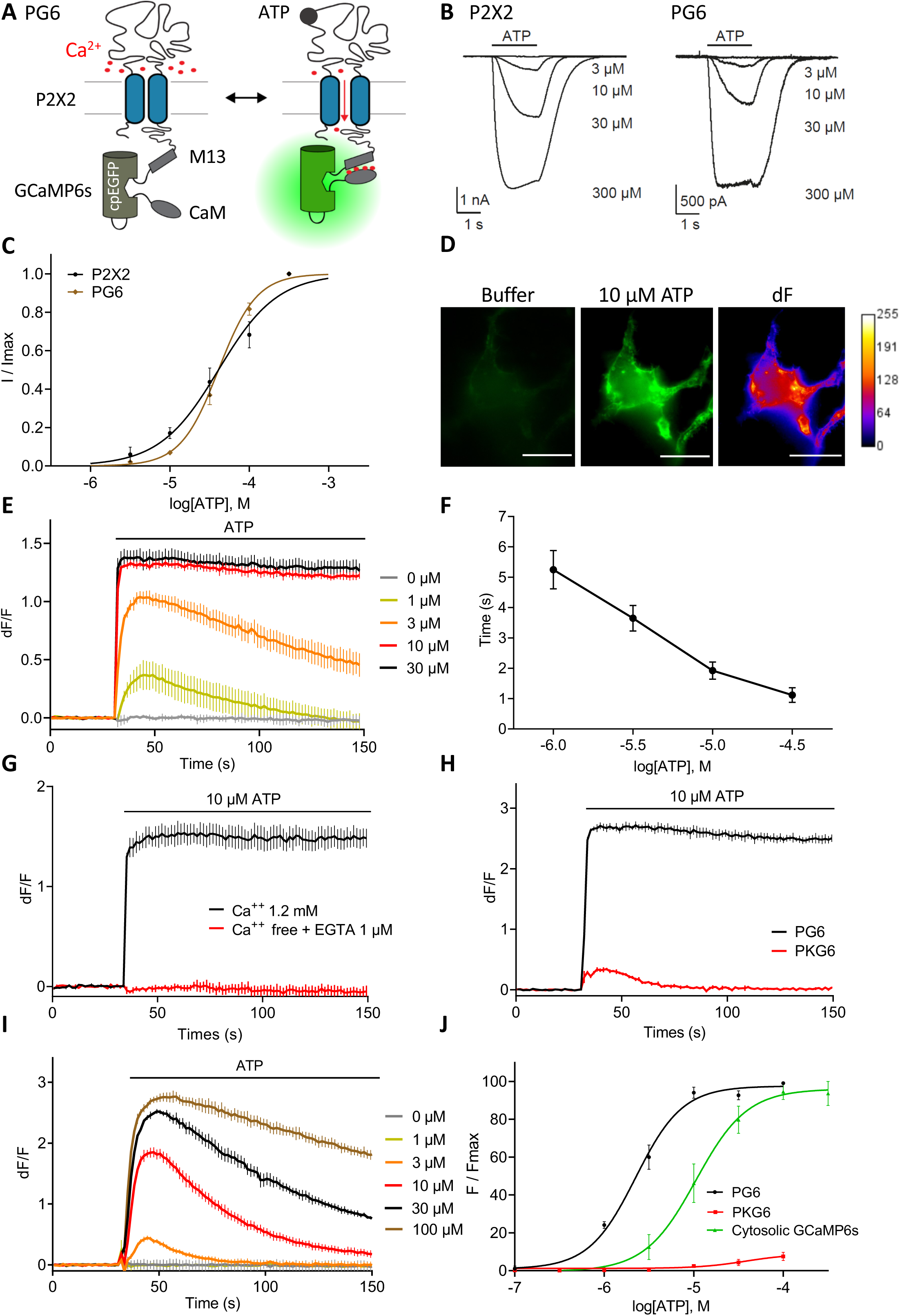
P2X2-GCaMP6s (PG6) fusion is functional and displays concentration dependent ATP-evoked fluorescence changes. **A.** Cartoon illustrating the biosensor and the change in GCaMP6s fluorescence during P2X2 activation. **B and C.** Representative traces (B) and normalized dose-response curves (C) of ATP evoked current measured by whole-cell recording in HEK cells expressing wild type P2X2 receptors or PG6 fusion. **D.** Representative images showing fluorescence changes (λexc/em 485/540) of a HEK cell expressing PG6 after ATP application (10 μM). Images show pseudo colored dF after F0 subtraction. Scale bar 20 μm. **E.** Example traces generated from a plate reader of changes in dF/F values triggered by increasing concentration of ATP in HEK cells expressing PG6. **F.** Relation between PG6 time constant of activation (τON) and ATP concentration. τON was calculated from N = 8 independent experiments as described in panel E. **G and H.** Representative traces from a plate reader of changes in dF/F values triggered by 10 μM ATP. HEK cells expressed either PG6 in presence or absence of extracellular calcium (G), or PG6 and PKG6 (H). **I.** Representative traces from a plate reader of changes in dF/F values triggered by increasing concentration of ATP in HEK cells expressing the cytosolic GCaMP6s. **J.** Normalized concentration-response curves for ATP evoked changes in F/Fmax at the PG6 and PKG6 fusions and cytosolic GCaMP6s. Curves were generated from N > 4 independent experiments. Data are expressed as means ± SEM in all panels. For panels E, G, H and I representative data from n = 3 wells per condition.

We also analyzed whether calcium influx through other ion channels localized at the plasma membrane could contribute to PG6 fluorescence. We first co-expressed in HEK cells PG6 with GluN1 and GluN2A subunits of the NMDA receptor, a ligand-gated channel with significant calcium permeability. In these cells, application of 50 μM NMDA and 10 μM glycine evoked a small fluorescence signal (20.3 ± 3.9 %, n = 3) of that evoked by 10 μM ATP (Fig. 2A and 2E). A similar signal was obtained in cells co-expressing PKG6 and GluN1/GluN2A subunits (16.7 ± 4.5 %, n = 3) (Fig. 2B and 2E). By comparison, in HEK cells co-expressing GluN1/GluN2A and the cytosolic GCaMP6s, both NMDA/glycine and ATP application evoked strong fluorescence signals (Sup Fig. 1D) indicating that calcium influx through NMDA channel marginally activates PG6. Similar experiments were performed by co-expressing TRPV1 and PG6. In this case, 100 nM capsaicin application induced a fluorescence signal of 31.9 ± 8.1 % of the fluorescence evoked by 10 μM ATP (Fig. 2C and 2F). In cells co-expressing TRPV1 and PKG6, capsaicin (100 nM) and ATP (10 μM) evoked fluorescence signals were 18.4 ± 5.3 % and 8.1 ± 2.9 % (Fig. 2D and 2F). Here also, in HEK cells co-expressing TRPV1 and the cytosolic GCaMP6s, both capsaicin and ATP application evoked strong fluorescence signals (Sup. Fig 1E). NMDA and capsaicin evoked fluorescence in PG6and PKG6 expressing cells were not significantly different, yet both signals were significantly smaller than fluorescence evoked by 10 μM ATP in PG6 expressing cells. Together these results support that PG6 fluorescence signal is mainly triggered by calcium influx through the pore of the channel and that any other sources of intracellular calcium increase minimally contribute to the fluorescence signal.

**Figure 2:**
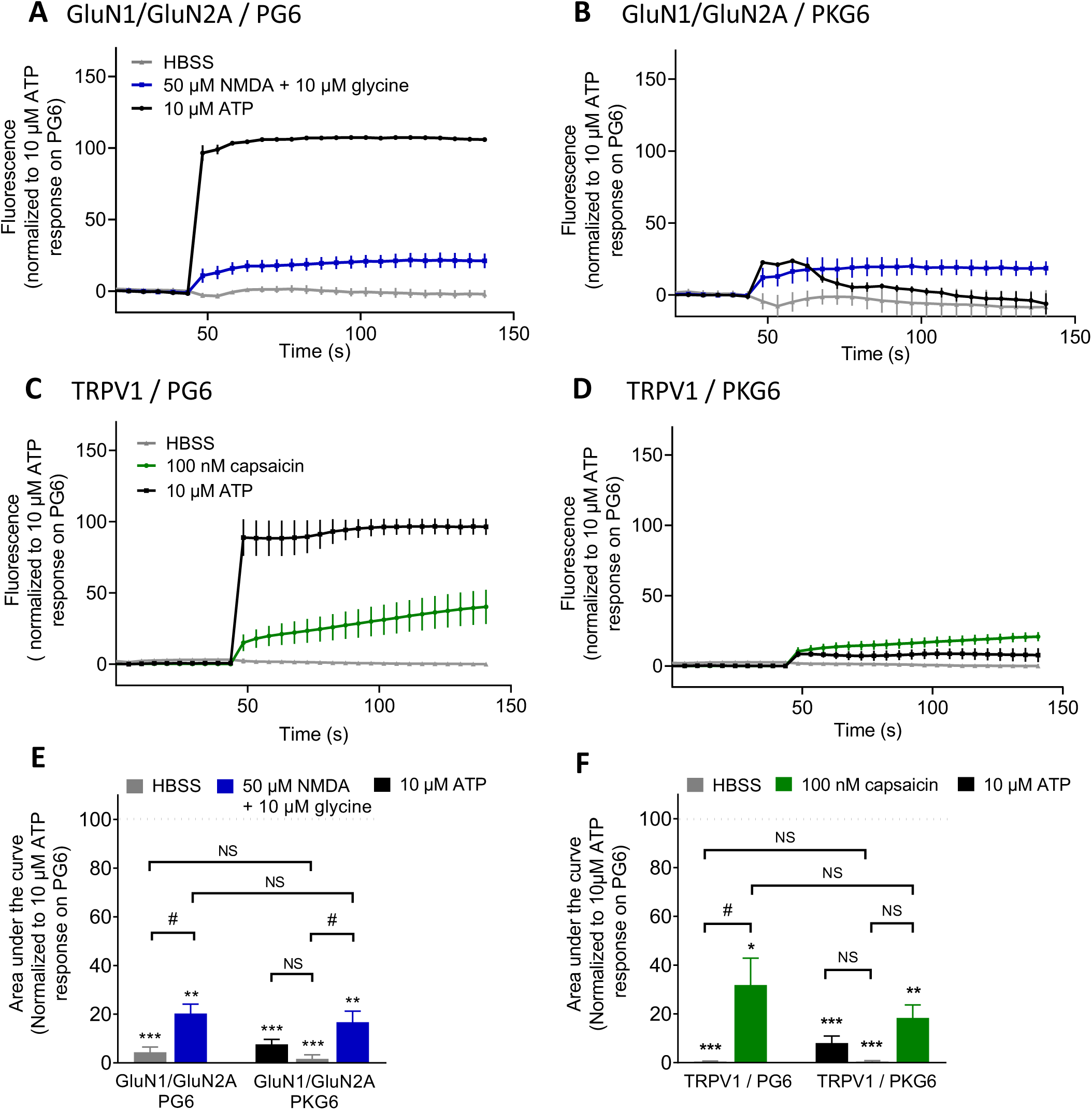
PG6 fluorescence signal is mainly triggered by calcium influx through the pore of the channel. **A and B.** Representative traces of normalized fluorescence changes evoked by ATP (10 μM, black), HBSS (gray) or NMDA/glycine (50/10 μM, blue) in HEK cells expressing NMDA receptor subunits GluN1 and GluN2A in combination with either PG6 (A) or PKG6 (B). Data were generated from a plate reader with n = 3 wells per condition. **C and D.** Representative traces of normalized fluorescence changes evoked by ATP (10 μM, black), HBSS (gray) or capsaicin (100 nM, green) in HEK cells expressing TRPV1 in combination either with PG6 (C) or PKG6 (D). Data were generated from a plate reader with n = 3 wells per condition. **E.** Plots of area under the fluorescence curve obtained from data shown in panels A and B. Values were normalized to response evoked by 10 μM ATP in PG6 expressing cells. **F.** Plot of area under the fluorescence curve obtained from data shown in panels C and D. Values were normalized to response evoked by 10 μM ATP in PG6 expressing cells. In all panels data are expressed as means ± SEM of N ≥ 3 independent experiments. * p< 0.05, ** p < 0.005, *** p < 0.001, one-sample t-test compared to 100% reference value. # p < 0.005, NS non-significant, one-way ANOVA with FDR corrected post-hoc tests.

The fusion strategy was extended to other P2X subunits, which were linked to GCaMP6s by C-terminal fusion. Among the six subunits individually expressed in HEK cells, dose-dependent ATP evoked fluorescence signals were observed for P2X4, P2X5 and P2X7 (Fig. 3A, 3B and 3C). Apparent affinities for ATP were 1.2 ± 0.3 μM, 8.1 ± 3.1 μM and 72.3 ± 28 μM, for P2X4, P2X5 and P2X7, respectively (Fig. 3E and 3F). In the case of P2X7, a higher apparent affinity of 5.2 ± 2.5 μM for BzATP was observed (Fig. 3C and 3E). No fluorescence signal was observed for P2X1 and P2X3 presumably because the rapid desensitization rates of these two receptors are faster than the acquisition frequency of the plate reader (1 Hz). Regarding P2X6, this subunit does not form functional homomeric channel (Fig. 2D). As previously shown for PG6, ATP-evoked fluorescence at P2X4-GCaMP6s was abolished in the absence of extracellular calcium (Sup. Fig. 1B) and mutation of ATP binding site (P2X4-K69A-GCaMP6s) significantly reduced ATP-evoked fluorescence, although ATP concentrations above 10 μM evoked fluorescence signals stronger than observed for PKG6 (Sup. Fig. 1A and 1C and Fig. 1H and 1J), presumably because internalized P2X4 receptors might be more sensitive to P2Y-evoked cytosolic calcium increase (Bobanovic et al., 2002). Activity of P2X2- and P2X4-GCaMP6s were also analyzed in the astrocytoma cell line 1321N1 that does not express endogenous P2Y receptors. 1321N1 cells were transduced using lentiviruses expressing either WT or K69A mutant of P2X2- or P2X4-GCaMP6s, and ATP-evoked fluorescence was recorded by video-microscopy at an acquisition rate of 1Hz (Sup. Fig. 2A, 2B and 2D). Ten seconds applications of increasing ATP concentrations induced dose-dependent fluorescence signal in cells transduced by WT receptors but not in cells expressing mutant receptors (Sup. Fig. 2C and 2E). These experiments indicate that the fluorescent signal of P2X-GCaMP6s accurately reports the activation of P2X2, validating the use of these new tools for medium to high throughput screening.

**Figure 3:**
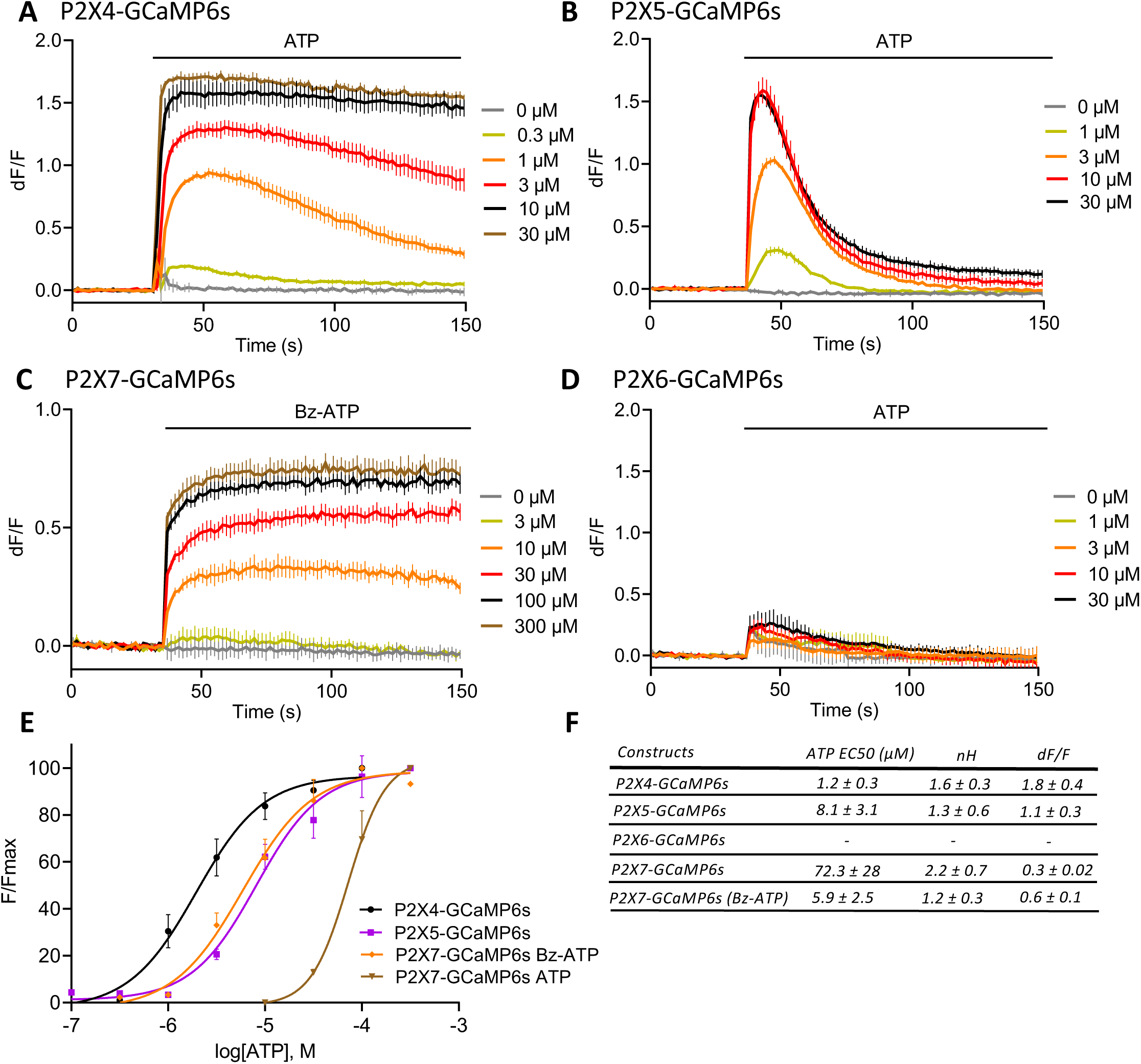
P2X-GCaMP6s fusion strategy extended to others P2X receptors. **A to D.** Representative traces of changes in dF/F values triggered by increasing concentration of ATP (A, B and D) or Bz-ATP (C) in HEK cells expressing P2X4-GCaMP6s (A), P2X5-GCaMP6s (B), P2X7-GCaMP6s (C) or P2X6-GCaMP6s (D). Data were generated from a plate reader with n ≥ 3 wells per condition. **E.** Normalized concentration-response curves for ATP or Bz-ATP evoked changes in F/Fmax at the P2X4-, P2X5-, P2X6- and P2X7-GCaMP6s fusions. Curves were generated from N = 3 independent experiments. Data are expressed as means ± SEM in all panels. **F.** Summary of EC50, Hill coefficient (nH) and maximal dF/F for P2X4-, P2X5-, P2X6- and P2X7-GCaMP6s fusions expressed in HEK cells.

We next asked whether ATP biosensors could be used to investigate ATP signaling in neurons. To that aim we generated lentiviruses expressing either PG6 or PKG6 under the control of the CaMKII alpha promoter to transduce primary culture of hippocampal neurons. Expression and subcellular localization of PG6 were analyzed by immunocytochemistry and western blotting. PG6 is present in neuronal cell body as well as in dendritic compartment as shown by the colocalization of GFP and MAP2 immunostaining (Fig. 4A). Both PG6 and PKG6 are expressed at the plasma membrane, as demonstrated by cell surface biotinylation and subsequent neutravidin pull down (Sup Fig. 3A). Further, subcellular fractionation reveals that PG6 and PKG6 are enriched in the synaptosomal fraction (Sup Fig. 3B and 3C).

**Figure 4:**
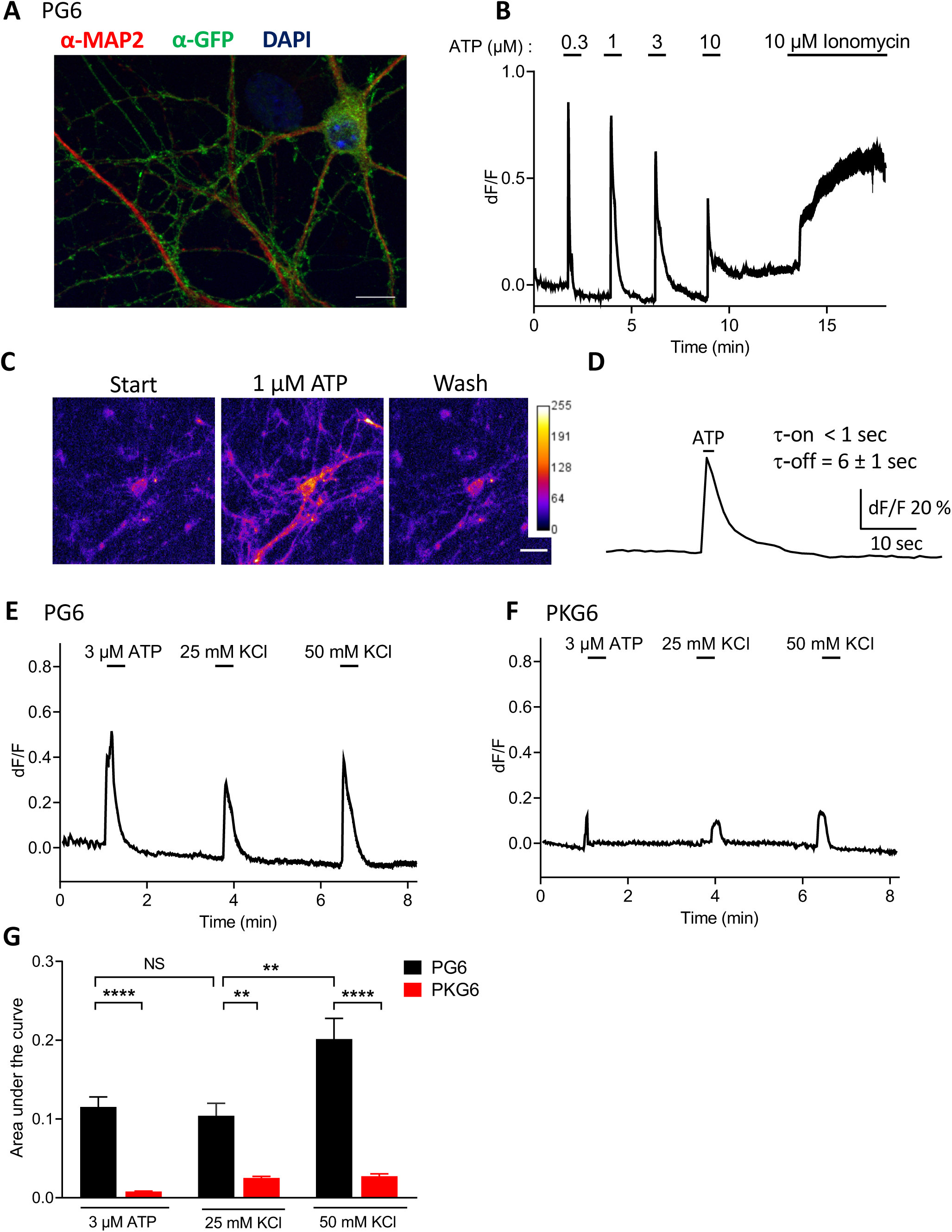
Expression and functional analysis of ATP sensors in hippocampal neurons. PG6 was virally expressed in neurons under the control of the CaMKII alpha promoter. **A.** transduced neurons were stained using anti-GFP (green) and MAP2 (red) antibodies and nucleus was stained with DAPI (blue). PG6 is localized in the whole dendritic tree as well as in varicosities. Scale bar 20 μm. **B.** ATP-evoked fluorescence in neurons expressing PG6 was recorded by video microscopy (λem/exc=485/538, acquisition rate 1 Hz). ATP was applied by gravity for 10 sec. Note the apparent desensitization of the response for high ATP concentrations. 10 μM ionomycin was applied for 5 min at the end of the experiment. N = 1 experiment, n > 10 neurons. **C.** Representative images of ATP-evoked fluorescence. Intensity of fluorescence was color coded. Scale bar 20 μm. **D.** Kinetics of ATP-evoked fluorescence. Representative recording showing fluorescence evoked by 0.3 μM ATP applied for 2 sec. τon and τoff were calculated with OriginPro 2017 using an exponential fit function. N = 2 independent experiments, n > 10 neurons **E.** Representative fluorescence recording of KCl-evoked activation of PG6 in neurons. 3 μM ATP was first applied as above followed by 25 mM and 50 mM KCl. Fluorescence was acquired as above. KCl-evoked depolarization induced ATP release which in turn activated PG6. **F.** Similar experiment using PKG6. Only a small and transient fluorescence signal were evoked by depolarization. For E and F, N = 2 independent experiments, n > 12 neurons. **G.** Plot of area under the curves obtained from panel E and F. One-way ANOVA with Tukey’s multiple comparisons test, **, *** and **** indicate p value < 0.01, < 0.001 and < 0.0001 respectively.

Functional analysis by video microscopy showed that in neurons, PG6 can be activated by sub-micromolar ATP concentration. Fluorescence signal in response to 300 nM shows rapid onset below 1 second but slower offset (6 sec) (Fig. 4B and 4D), which is likely due to a slow washout of the agonist. At 1 μM ATP, fluorescence signals show fast activation and rapid return to base line while at 10 μM ATP and above, fluorescence remains high even after ATP wash out (Fig. 4B and supplementary Movie 1). This may suggest that in these conditions and in the absence of any inhibitor or antagonist of synaptic transmission, depolarization evoked by sustained P2X2 activation can elicit additional calcium influx, contributing to GCaMP6s fluorescence. We reasoned that KCl-evoked neuronal depolarization should elicit endogenous ATP release that can possibly be detected by PG6 sensor. Hippocampal neuronal cultures were transduced as above either with PG6 or PKG6. Functional expression of the WT sensor was first tested by 10 seconds application of 3 μM ATP followed by successive 10 seconds applications of 25 and 50 mM KCl. In neurons expressing PG6, KCl-evoked depolarization induced transient fluorescence increase which intensity depends on the concentration of KCl (Fig. 4E). These increases of fluorescence were specifically due to direct ATP activation of the sensor since in neurons expressing PKG6, KCl applications evoked only minimal changes of fluorescence (Fig. 4F). As shown on Fig. 4G, the changes in fluorescence evoked by 25 mM KCl or by 3 μM ATP applications were not statistically different, suggesting that neuronal depolarization results in ATP release that can reach few micromolar at the plasma membrane. These results indicate that PG6 acts as an ATP biosensor with a sensitivity in the low micromolar range.

We thought to improve the sensitivity of the PG6 biosensor by introducing point mutations which enhance ATP potency for P2X2 receptor (Li et al., 2004). Five point mutations were individually introduced in PG6: I40A, I328A, P329A, N333A and V343A, and ATP potency for these different mutants were compared to that of the WT sensor (Fig. 5A, 5B and 5C). Out of these five mutants, ATP displayed a significant gain of potency for three of them: I328A, P329A and N333A with EC50s of 0.204 ± 0.03, 0.278 ± 0.07 and 0.327 ± 0.04 μM, respectively. Maximal ATP-evoked fluorescence intensities of these mutants were different (Fig. 5B). Normalization of fluorescence intensities to that of PG6 shows that maximal intensities of N333A and I40A were close to that of WT, while that of P329A was intermediate at 69.9 %, (95 % confidence interval (CI) 61.3 to 81.1), and I328A and V343A clearly reduced at 34.9 % (CI 30 to 41.5) and 29.7 % (CI 26.7 to 33), respectively. Only P2X2-N333A-GCaMP6s (stated below as PNG6) and P2X2-P329A-GCaMP6s (PPG6) were further considered. As illustrated in Fig. 5D and 5E, application of 300 nM ATP triggers strong fluorescence signals in cells expressing PNG6 and PPG6 (dF/F = 0.43 ± 0.04 and 0.33 ± 0.08, respectively) but not in cells expressing PG6. Functional properties of PNG6 and PPG6 were also investigated by whole cell recording in HEK cells. Compare to WT sensor, ATP activated currents at PNG6 and PPG6 with similar activation kinetics but somehow apparent slower deactivation (Sup Fig. 4A). Here also the two mutants displayed gain in apparent affinity for ATP; EC50s were 45.2 ± 7.8 μM, 7.9 ± 0.4 μM and 4.5 ± 0.3 μM for PG6, PNG6 and PPG6, respectively (Sup Fig. 4B).

**Figure 5:**
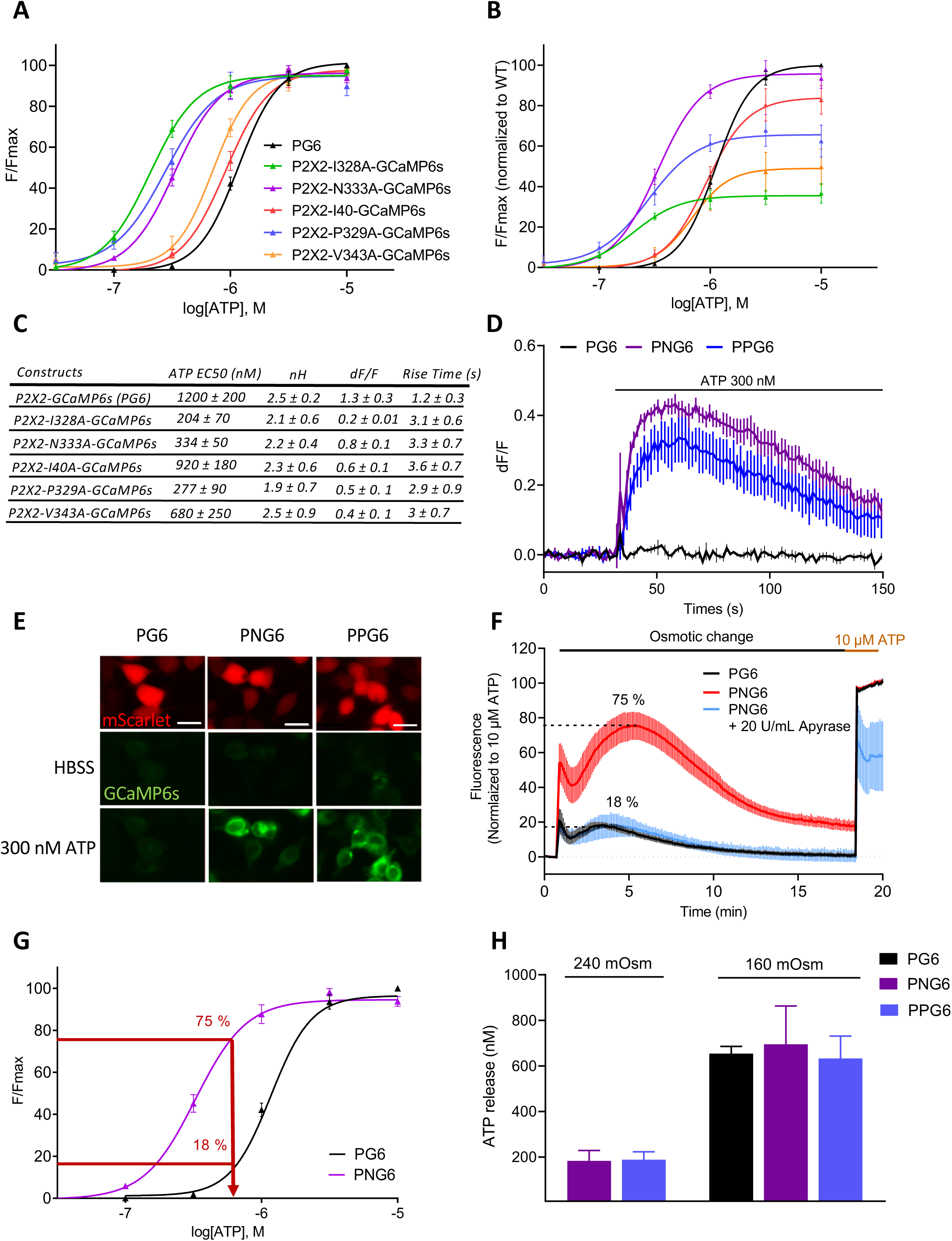
Functional characterization of high sensitivity ATP sensors. **A.** ATP potency at different single residue mutated PG6. Dose-response curves for ATP were performed in HEK cells transfected with each mutant. Normalized curves were generated from N = 4 independent experiments. **B.** Dose-response curve for each mutant were normalized to that of PG6, showing that some mutations have significant lower response to saturating dose of ATP. N = 4 experiments. **C.** Table of the main characteristics for each mutant. **D.** Example traces generated from a plate reader of changes in dF/F values triggered by 300 nM ATP in HEK cells expressing PG6, PNG6 (N333A mutant) and PPG6 (P339A mutant); n = 3 wells **E.** Epifluorescence images showing fluorescence changes (λexc/em 485/540) of HEK cells expressing PG6-P2A-Scarlet, PNG6-P2A-Scarlet and PPG6-P2A-Scarlet during 300 nM ATP application. Scale bar 20 μM. **F.** Sensing ATP release from HEK cell during hypo-osmotic stimulation. Comparison of fluorescence signals recorded in a plate reader from cells expressing PG6 or PNG6 during a 160 mOsm application. Data are normalized to the fluorescence evoked by a 10 μM ATP application. 20 U/mL Apyrase strongly inhibited hypotonicity-evoked fluorescence in cells expressing the high sensitive ATP sensor; n = 3 wells **G.** Estimation of the maximal concentration of ATP release during hypotonic challenge. Maximal fluorescent values measured from cells expressing PG6 or PNG6 and reported on their respective dose-response fitted curve. **H.** Group data quantification of ATP release during 240 and 160 mOsm hypotonic challenge of cells expressing PG6, PNG6 and PPG6. N = 6 independent experiments. Data from all panels are mean ± SEM.

We finally evaluated to what extend these different sensors could detect endogenous ATP release. Hypotonicity is a known stimulus triggering ATP release from virtually all cells (Okada et al., 2018). We tested in HEK cells whether ATP release induced by a reduction of extracellular osmolarity could be detected by PNG6 and PPG6 biosensors. A 160 mOsm saline solution (50 % reduction of osmolarity) was applied to cells expressing either PG6 or the more sensitive PNG6 biosensors, and fluorescence variations were recorded prior a final application of 10 μM ATP (Fig. 5F). In cells expressing PNG6 sensor, a biphasic increase of fluorescence was observed. The first transient peak is likely due to ATP release evoked by the mechanical stimulation induced by the solution change. The second phase developed slowly with a maximum reached around 5 minutes after the reduction of osmolarity, and then slowly decreased to reach a plateau. This second phase likely corresponds to the development of cellular regulatory volume decrease, during which ATP is being released. Maximum of fluorescence represented 75.4 ± 8.1 % of the signal evoked by a 10 μM ATP stimulation. In the presence of 20 U/mL apyrase, a soluble ectonucleotidase, hypotonicity-evoked fluorescence variations were strongly inhibited, further supporting a hypotonicity-evoked ATP release. In cells expressing PG6 sensor, a similar biphasic fluorescence response was observed, albeit with a much lower intensity, 18.2 ± 1.8 % of the 10 μM ATP-evoked response. Based on these values, we estimated the maximal ATP concentration reached during a hypotonic environment by extrapolating the maximal fluorescence values measured of cells expressing either PNG6 or PG6 in hypotonic conditions to their respective fitted dose-response curves. In both cases similar estimations were obtained with extracellular ATP concentration of 695 ± 137 nM and 656 ± 26 nM for the PNG6 and PG6 sensors, respectively (Fig. 5F, 5G and 5H). When similar experiments were performed using a 25 % reduction of osmolarity (240 mOsm), no change in fluorescence was observed in cell expressing PG6 sensor (data not shown), whereas a 18 ± 3.8 % increase of fluorescence was detected in cells expressing PNG6 biosensor (Sup. Fig. 5A); leading to an estimated ATP concentration of 183 ± 32 nM (Fig. 5H). Essentially identical results were obtained with the PPG6 sensor (Fig. 5H, sup. Fig. 5B). For comparison, extracellular ATP was quantified every 5 min in the same condition of reduced osmolarity using luciferase based ATP quantification kit. A similar slowly developing, bell shaped ATP increase was observed, with a maximum reached 5 to 10 minutes after the start of the osmotic challenge (Sup. Fig. 5C).

Red shifted P2X2 and P2X2-N333A sensors were also generated by fusion of the respective cDNA to either RCaMP2 or jRGECO1a (Sup. Fig.6). P2X2-RCaMP2 and P2X2-N333A-RCaMP2 show the same pharmacology as previously described for the GCaMP6s version, while to the difference with PKG6, P2X2-K69A-RCaMP2 was not activated by ATP up to 50 μM (Sup. Fig. 6A). ATP EC50s for P2X2-RCamp2, P2X2-N333A-RCaMP2 and P2X2-N333A-jRGECO1a were respectively 2.6 ± 0.4 μM, 448 ± 77 nM and 358 ± 41 nM (Sup. Fig. 6A, 6D and 6F). We next compared fluorescence evoked by ATP for P2X2-N333A-RCaMP2 and P2X2-N333A-jRGECO1a. At 1 μM ATP, dF/F signals were twice larger for P2X2-N333A-jRGECO1a compared to P2X2-N333A-RCaMP2 (dF/F = 0.63 ± 0.03 and 0.35 ± 0.01, n = 3 wells, respectively) (Sup. Fig. 6B). Similarly, P2X2-N333A fused to either RCaMP2 or jRGECO1a dynamically detected ATP release during hypotonic challenge with fluorescence values equal to 80.3 ± 3.5 % (RCaMP2) and 70.7 ± 5 % (jRGECO1a) of the response evoked by 10 μM ATP (Sup. Fig 6C). Estimation of the maximal ATP concentration released during the hypotonic challenge, 650 ± 58 nM and 743 ± 24 nM (N=3) for P2X2-N333A-RCaMP2 and P2X2-N333A-jRGECO1a, respectively (Sup. Fig. 6D and 6E), are close to those observed with PNG6 and PPG6 sensors. These experiments indicate that red shifted ATP sensors have the same sensitivity and properties as the green versions and support that both green and red versions can be used in combination to track the dynamic of extracellular ATP release in two different cell types within a cellular network.

## Discussion

In this study we engineered P2X receptors with the aim i) to directly track their activity and ii) to detect extracellular ATP with fast kinetics and high sensitivity. By fusing the genetically encoded calcium indicator (GECI) GCaMP6s to the carboxy terminal end of P2X subunits, we generated versatile probes which allow both monitoring P2X receptor activity and biosensing extracellular ATP release.

The principle of tracking P2X receptor activity by the mean of a C-terminal fusion with a FRET-based GECI was previously described (Richler et al., 2008). Although this approach demonstrated its specificity *in vitro* and *in vivo*, the use of Cameleon YC3.1 as a fusion partner presented several limitations among which rather slow kinetics and a difficult implementation. We thought that with the improvement of single wavelength GECI based on circularly permuted fluorescent proteins (GCaMP) most of these limits could be overcome.

Although most of our experiments were based on the P2X2 biosensors, GCaMP6s fusions also report the activity of three additional P2X receptors (P2X4, P2X5 and P2X7) out of the seven possible, which all display ATP potencies, and BzATP in the case of P2X7, in agreement with published values (Jarvis and Khakh, 2009). In our experimental conditions, we were not able to track P2X1 and P2X3 activity as these two receptors desensitize with sub second rates (North and Surprenant, 2000), which is too fast to be detected in our multi well fluorescent plate reader assay. For these subunits, as well for P2X6 that does not form functional receptors, further experiments using heteromeric receptor expression and/or high-speed drug application combined with high rate of fluorescence acquisition should help to extend our observations to all P2X subunits.

One potential bias of using GCaMP6s as a reporter of P2X receptor activity might come from fluorescence evoked by intracellular calcium increase unrelated to P2X receptor permeability. Our results using P2X2-K69A, a mutant unable to be activated by ATP (Jiang et al., 2000), support that both in HEK cells and neurons, once tethered to P2X2 C-Terminus, GCaMP6s is almost exclusively activated by calcium influx through P2X permeation pathway. Even though we cannot totally exclude that in some conditions a P2X-independent activation of GCaMP6s might occur, experiments of co-expression of PKG6 with highly calcium permeant channels such as NMDA receptor or TRPV1 clearly indicate that calcium influx through other calcium permeant channels minimally activate PG6. Systematic control experiments using PKG6 should allow to estimate potential contamination of the P2X-GCaMP6s signal. Alternatively, assuming that local intracellular calcium concentration is the highest at the vicinity of the inner mouth of the P2X permeation pathway, using GCaMPs in which point mutations are introduced in EF hand and RS20 peptide in order to lower the affinity for calcium (Helassa et al., 2016) should reduce nonspecific activation of PG6. Such a strategy was successfully used in the case of GenEPi, a Piezo1-based fluorescent reporter for visualizing mechanical stimuli (Yaganoglu et al., 2019).

A clear difference in ATP potency on PG6 was observed whether EC50s were measured by whole cell recording or by acquisition of GCaMP6s fluorescence. Such a difference was not previously reported for the P2X2-Cameleon. Yet, the affinities of calcium sensing modules and the conformational changes required to generate fluorescence of GCaMP6s and Cameleon present clear differences (Chen et al., 2013; Miyawaki et al., 1999). It is thus possible that GCaMP6s can report with more accuracy discrete channel opening than Cameleon. We also noticed that ATP potency varies as a function of the cell type in which P2X-GCaMP6s are expressed, particularly in neurons in which sub-micromolar ATP evokes significant activation of P2X2-GCaMP6s compared to HEK cells. These differences in potency might be due to the lipid composition of the plasma membrane, e.g. in cholesterol, which is known to modulate certain P2X receptors (Murrell-Lagnado, 2017). Alternatively, in neurons proteins associated with P2X2 might also modulate apparent affinity of the channel for ATP (Chaumont et al., 2008). In our case, electrophysiological and fluorescence measurements were realized on two HEK cell lines maintained in different laboratory, which can account for the observed differences in EC50s.

When expressed in neurons, PG6 is localized at the plasma membrane and distributed in the dendritic tree. Purification of synaptoneurosome fraction also supports its enrichment at synapses. Although we did not precisely determine a post versus pre-synaptic localization of PG6, the dendritic expression strongly suggests a post synaptic localization of the protein. This is in agreement with previous studies showing that in neurons P2X2 traffics to dendrites and to the postsynaptic element where it likely modulates glutamatergic transmission (Emerit et al., 2016; Pougnet et al., 2014; Richler et al., 2011). In neurons, application of a sub micromolar ATP concentration evokes transient fluorescence changes, while higher concentrations trigger both transient and plateau response. It is difficult to estimate whether fluorescence signal is mediated solely by calcium flowing through P2X2 channel or whether P2X2-evoked depolarization might activate voltage gated calcium channels and subsequent calcium entry, indirectly contributing to GCaMP6s activation. We observed that KCl-evoked neuronal depolarization leads to an activation of PG6 sensor in dendrites, which likely results from synaptically released ATP. Yet KCl-evoked depolarization did not trigger PKG6 fluorescence. This supports that calcium-entry through voltage-gated calcium channels minimally contributes to activation of GCaMP6s tethered to P2X2 receptor. Altogether, these results suggest that in primary neuronal cultures, the PG6 biosensor is capable of detecting endogenous ATP release following chemical depolarization. Future experiments will be necessary to further demonstrate that PG6 biosensor can efficiently detect ATP release in response to neuronal activity in acute slice and eventually *in vivo*.

Because extracellular ATP concentrations are usually low (Beigi et al., 1999; Gourine et al., 2005), sensitivity of ATP sensors is an important parameter. The introduction of point mutation in P2X2 (e.g. N333A, P329A) known to improve the potency of the receptor for ATP (Li et al., 2004), leads to the generation of extracellular ATP biosensors with a sensitivity covering a 100 to 1 000 nM range. These sensors dynamically detected ATP release induced by a reduction of extracellular osmolarity, which we estimated to reach 600 nM. Although these later experiments were performed in a plate reader and on cell population, we provide evidence that these sensors can detect ATP release at single cell level in 1321N1 and neurons. Altogether, these data provide proof of concept that PG6 could be instrumental to dynamically detect events of ATP release in diverse cellular contexts.

Different optical, genetically engineered, extracellular ATP biosensors have been generated, which all display different specificities on four main parameters i.e. sensitivity, kinetics, brightness and dynamic range (Wu and Li, 2020). Although we did not perform in depth characterization of these parameters, P2X-GCaMP6s sensors characteristics outperform that of most other optical biosensors. First, P2X-GCaMP6s sensors offer a large range of sensitivity from nanomolar range (PNG6) up to millimolar range (P2X7-GCaMP6s). Second, activation kinetics of PG6 appear very fast (< 1 sec), particularly in neurons, even though our analysis remains constrained by a 1-2Hz acquisition rate. This is not surprising since both P2X2 and GCaMP6s activates within millisecond time scale (Chen et al., 2013; Markwardt, 2007). Off rate kinetics of PG6 fluorescence are slower, yet this parameter is constrained by the inactivation kinetics of GCaMP6s which is in the range of several seconds. Yet, these characteristics of PG6 can be further improved through the use of ultrafast version of GCaMP6 (Helassa et al., 2016). Although PG6 kinetics presented here are rough estimations, PG6 is likely to present the fastest kinetics among the all existing optical ATP biosensors. Finally, owing to the dynamic range performance of GCaMP6 series, P2X-GCaMP6s sensors present very good dF/F0, greater than one for most constructs expressed in HEK cells. Finally, we also generated red-shifted variants of P2X-based ATP sensors. Particularly, P2X2-jRGECO1a series present a dF/F close to that of PG6. These two colors ATP sensors should allow to get a better understanding of the ATP signaling dynamic, particularly in the brain where paracrine action of ATP is thought to contribute to network activities in both physiological and pathological conditions. Altogether, performances of P2X-GCaMP6s series should allow to dynamically detect extracellular ATP release events with high sensitivity and rapid kinetics, although the activity of P2X receptors might represent a limitation to its use in specific conditions. These sensors should therefor represent powerful tools to investigate mechanisms of ATP release and signaling as well as to track P2X receptor activity *in situ*.

## Materials and Methods

### cDNA cloning and site-directed mutagenesis

The GCaMP6s cDNA (gift from Douglas Kim & GENIE Project, Addgene plasmid # 40753) was subcloned in pWPT-EF1α-IRES-DsRed2 or pWPT-EF1α-P2A-mScarlet lentiviral vectors by enzymatic digestion. P2A-Scarlet (Bindels et al., 2017) was produced by gene synthesis (Eurofins Genomics) and directly subcloned in appropriate expression vector. For rat P2Xs-GCaMP6s fusions, GCaMP6s was PCR inserted in frame at the C-terminal tail of P2X subunits. P2X2-GCaMP6s cDNA was inserted in pWPT-EF1α-IRES-DsRed2, pWPT-EF1α-P2A-mScarlet or pWPT-CaMKIIα lentiviral vectors by enzymatic digestion. A similar strategy was used to fuse P2X2 subunits with RCaMP2 (Inoue et al., 2015) (gift from Dr. Perroy, Institut de Genomique Fonctionnelle, CNRS, Montpellier, France) and jRGECO1a (gift from Douglas Kim & GENIE Project, Addgene plasmid # 61563), or other P2X subunits to GCaMP6s. Mutations were generated in the rat P2X2 receptor using the Q5 Site-Directed Mutagenesis Kit (NEB). Each mutation was verified by DNA sequencing and subcloned in pWPT-EF1α-P2X2-GCaMP6s-IRES-DsRed2 or pWPT-EF1α-P2X2-GCaMP6s-P2A-mScarlet lentiviral vectors.

### HEK-293T cells culture

HEK-293T cells (ATCC CRL-3216) were maintained in Dulbecco’s modified Eagle’s medium (DMEM-GlutaMax, Thermofisher), 10 % fetal bovine serum (FBS, Thermofisher) and 1 % Penicillin-Streptomycin (PS, 10000 U/mL, Thermofisher). Cells were grown in a humidified atmosphere of 95 % air/ 5 % CO2 at 37°C in a cell culture incubator. Cells were split 1/10 when the confluence reached 90 %, generally every 3 to 4 days.

### HEK-293T cell transfections

For GCaMP6s fluorescence recording, HEK-293T cells were plated in six-well plates 1 day before transfection. Cells were transfected, depending on the plasmid, with 0.1 to 1 μg of plasmid for each well using Lipofectamine 2000 (Thermofisher) according to the manufacturer’s recommendation. 24 hours after transfection, cells were split in poly-L-ornithine (Sigma) coated 96-well plate and incubated at 37°C for 24h or 48h before imaging experiments. For co-transfections experiments, 0.5-1 μg of plasmids were used for PG6, PKG6 and GCaMP6s, 60 ng for TRPV1 and GluN2A and20 ng for GluN1. In experiments where GluN1/GluN2A were co-expressed, cells were maintained in 10 μM D-AP5 (Tocris) to ovoid cellular toxicity.

For patch clamp experiments, trypsin treated HEK cells were seeded onto poly-Lysine (Sigma) pretreated glass coverslips in 35-mm dishes 1 day before transfection and incubated at 37°C with 5 % CO2. Transfections were carried out using calcium phosphate precipitation using 0.1– 0.3 μg of P2X2 or P2X-GCaMP6s plasmids. For P2X2 experiments, 0.3 μg of pCDNA3-GFP plasmid was added to identify putative co-expressing cells. Medium was changed 1 day after transfection and used within 24h.

### Lentivirus production

Versene treated HEK cells were seeded in 15 cm plate at 60 % confluence. 1 day later, cells were transfected with a combination of three plasmids: 5 μg of pMD2.G (gift from Didier Trono, Addgene plasmids #12259), 15 μg of psPAX2 (gift from Didier Trono, Addgene plasmids #12260) and 20 μg of the transfer plasmid. Transfections were carried out using calcium phosphate precipitation. 6 hours after transfection, HEK cells were washed with fresh medium (DMEM GlutaMax + 1 % FBS + 1 % P/S). 72 hours after transfection, the supernatant was collected and filtered (0.45 μm filter) to remove cellular debris. 40 % polyethylene glycol solution (PEG6000, Sigma) 4X was added to the supernatant and incubated overnight at 4°C and centrifugated for 30 min at 2600 g at 4°C. The pellet containing the lentiviral particles was resuspended in 100 μL PBS, aliquoted and stored at −80°C.

### 1321N1 Astrocytoma cell line and transductions

1321N1 cells were grown and cultivated in the same condition as HEK-293T cells. For lentiviral transduction cells were seeded in 6-well plates at 50 % confluence. 1 day later, cells were transduced with 6 μL of virus mixed with 6 μL of LentiBlast solution (3 μL of LentiBlast A solution and 3 μL of LentiBlast B solution, Ozbiosciences). 2-3 days after transduction, most cells expressed the transgene in a stable manner and were cultivated in DMEM + 10 % FBS + 1 % PS. For imaging experiments, cells were seeded at 50 % confluence in 35-mm dishes (Ibidi) 1day before experiment.

### Electrophysiological recordings

Currents were recorded using the whole-cell configuration of the patch clamp technique only from fluorescent cells. Cells were maintained at a holding potential of −60 mV. Patch pipettes (3-5 megaohms) contained (in mM) 140 KCl, 5 MgCl_2_, 5 EGTA, 10 HEPES, pH 7.3 adjusted with NaOH, (maintained at 300 mOsm). External solution contained (in mM) 140 NaCl, 2.8 KCl, 2 CaCl_2_, 2 MgCl_2_, 10 glucose, 10 HEPES, pH 7.3 adjusted with NaOH, (maintained at 300 mOsm), containing or not ATP, and was delivered through three parallel tubes placed immediately above the cell. These tubes are displaced horizontally with the aid of a computer-driven system (SF 77A Perfusion fast step, Warner) that ensures solution exchange in 5–10 ms. ATP (sodium salt, Sigma) was applied briefly (2 seconds), and for high ATP concentrations, pH was carefully adjusted with NaOH.

### Neuronal primary culture and transductions

Detailed protocol of the culture has been already described (Moutin et al., 2020). Hippocampi from newborn C57Bl/6J mice (P0 to P1) were collected in Hibernate-A medium (Thermofisher) + 1 % P/S, digested using 0.1 % papain at 37°C for 10 min and for 5 additional minutes in presence of DNAse1 (1 mg/ml, Roche). After stopping papain activity by adding MC+ media, a Neurobasal-A based medium (Thermofisher) supplemented with 2 % B27 (Thermofisher), 0.25 % Glutamax (Thermofisher), 0.5 mM L-glutamine (Thermofisher), 1 % P/S, 10 % FBS the tissue was mechanically dissociated using p1000 pipette. Dissociated tissue was centrifugated for 7 min at 300 g, cells were resuspended in MC+ media and were seeded in laminin and poly-L-ornithine pre-coated dishes (Ibidi) and incubated at 37°C. At DIV2, 1 μM AraC was added to reduce glial cell proliferation. At DIV3 the media was replaced by a BrainPhys based media (Stemcell technology) supplemented with 2 % B27, 0.25 % Glutamax, 1 % P/S. At DIV7, Cultures were transduced using 6 μl of lentiviruses for a 35 mm dish.

### Synaptosomes, plasma membrane biotinylation and western blotting

For synaptosomes experiments, transduced neurons were washed twice with PBS supplemented with 1 mM CaCl_2_ and 0.5 mM MgCl_2_ (PBS-CM) and lysed in Syn-PER reagent (Thermofisher) mixed with protease inhibitor cocktail (Thermofisher). Lysates were centrifugated for 10 min at 1200 g at 4°C. Supernatant was transferred in a new tube and 30 μl was kept as the homogenate fraction. The remaining supernatant was centrifugated for 20 min at 15000 g at 4°C. 30 μl of the supernatant was kept as the cytosolic fraction. The pellet was resuspended in 30 μl Syn-PER reagent and was kept as the synaptic fraction. Samples were diluted in LDS-Page 4X denaturant buffer (Invitrogen) supplemented with 5 % β-mercaptoethanol and heated for 5 min at 80°C. For biotynilation experiments, neurons were washed twice with PBS-CM and labeled using 1 mg/ml sulfo-NHS-LC-Biotin (Pierce) for 30 min at 4°C in PBS-CM. Cells were washed three times in PBS-CM supplemented with 10 mM Tris pH 7.4 to quench free reactive biotin. Cell lysis was performed for 30 min at 4°C under agitation in cell lysis buffer (100 mM NaCl, 20 mM HEPES pH 7.4, 5 mM EDTA, 1 % Triton X-100) mixed with protease inhibitors. Lysates were centrifuged (13.000 g, 10 min, 4°C) and the supernatants collected. Biotinylated proteins were purified with Neutradivin coated magnetic beads (Spherotech) for 1 hour at room temperature. After 3 washes with cell lysis buffer + protease inhibitor, proteins were eluted in LDS-Page 4X denaturant buffer (Invitrogen) supplemented with 5 % β-mercaptoethanol and heated for 5 min at 80°C. Proteins were resolved on gradient 4-12% Nu-PAGE gels (Thermofisher) in MOPS buffer (Thermofisher) and then transferred to a nitrocellulose membrane with the iBlot system (Thermofisher). Membranes were blocked with 0.1 % Tween20 in PBS (PBST) + 5 % non-fat milk for 1 hour at room temperature before incubating with primary antibody: anti-GFP (1:1000, Biolabs) or anti-actin (1:5000, DSHB) overnight at 4°C. Membranes were washed and incubated with the appropriate HRP-conjugated secondary antibody (1:10.000) for 2 hours at room temperature, then rinsed in PBS-T and revealed with SuperSignal West Pico substrate (Thermofisher). Signals were acquired with a Chemidoch touch (Biorad) and analyzed with ImageLab.

### Immunocytochemistry

At DIV15-DIV18, neurons on coverslips were washed twice with PBS, fixed with 4 % formaldehyde in PBS for 5 min at room temperature, and then washed three times with PBS for 5 min. Cells were blocked and permeabilized in PBS solution supplemented with 3 % bovine serum albumin and 0.1 % Triton X-100 (PBS-Perm) for 2 hours at room temperature. After overnight incubation at 4°C in PBS-Perm with rabbit anti-GFP (1:1000, Biolabs) and mouse anti-MAP2 (1:2000, Sigma) cells were washed and incubated with appropriate fluorescence-conjugated secondary antibodies (1:1000, Molecular Probes) for 2 hours at room temperature, then rinsed in PBS-Perm and incubated for 2 min with 300 nM of DAPI diluted in H_2_O. Coverslips were mounted on slides with a fluorescent mounting medium (DAKO). Images were acquired with an Axio Imager Z1 Apotome microscope (ZEISS) with 60 x objective lens (Z-stack with Z spacing of 1 μM). Raw images were treated using ZEN software and maximum intensity projections of these images are shown in the results.

### Plate reader recording

Transfected HEK cells seeded in 96-well plates were washed twice and incubated in FLEX buffer (HBSS (Thermofisher): 20 mM HEPES pH 7.4, 1 mM MgSO_4_, 3 mM Na_2_CO_3_, supplemented with 1.3 mM CaCl_2_) for 15 min at 37°C. For dose-response and pharmacological experiments, 2X concentrated drugs in FLEX buffer were delivered to the well. For experiments without extracellular calcium, cells were incubated for 15 min in the FLEX buffer without CaCl_2_ + 1 mM EGTA (Thermofisher). For hypotonicity experiments, H_2_0 Milli-Q water + 1.3 mM CaCl_2_ were added to the well to obtain 240 mOsm or 160 mOsm solutions. The osmotic concentration of each solution was verified with an osmometer. Fluorescence (λexc/em 485/540) was acquired with either a FlexStation 3 (Molecular Devices) equipped with automatic injectors or with an INFINITY 500 (TECAN) plate reader. Recording was performed at 1 or 0.15 Hz for the Flexstation 3 and the INFINITY 500, respectively. All the experiments were performed at 37°C.

### Time-lapse Imaging

For 1321N1 cells, videomicroscopy experiments were performed in FLEX buffer (see above) at least 10 days after transduction. For primary cultured neurons, cells were used at DIV15-DIV 18. Neurons were washed and incubated in ACSF buffer (150 mM NaCl, 2 mM CaCl_2_, 3 mM KCl, 10 mM HEPES pH 7.4, 10 mM D-Glucose). ACSF osmolarity was carefully adjusted to 320 mOsm using NaCl. External solution was delivered through a perfusion system allowing fast and local drug application. ATP (sodium salt, Sigma) and other compounds were applied briefly (10 sec) except for ionomycin which was applied for several minutes. All cells were imaged using an epifluorescence microscope (OLYMPUS IX70) equipped with an Evolve photometrics camera with a 20X water-immersion objective lens. Acquisitions were performed at 1 Hz (λexc/em 485/540). The microscope was driven by Metamorph software and data were analyzed with ImageJ. For images analysis, an ROI based analysis was performed with Image J. For 1321N1 cells, each cell was analyzed as an independent ROI. For primary neurons, each value is an average of 2-3 ROI localized in the dendrites and 1 ROI localized in the cellular body. τon and τoff values were calculated using an exponential fit function in the OriginPro software.

### ATP dosage by bioluminescence

ATP was detected by bioluminescence with the CellTiter-Glo 2.0 ATP kit (Promega) based on the manufacturer’s recommendation. Briefly, HEK cells were seeded in 24 well plate at a density of 2.5 10^5^ per well. Experiments were performed 24 hours later. After a 50 % osmotic choc (160 mOsm), an equivalent volume of the CellTiter-Glo reagent and cell supernatant were mixed in a 96 well plate, incubated for 5 min at room temperature and shake for 2 min at 37°C. The bioluminescence was quantified with a plate reader (INFINITY 500). In parallel, the quantity of cells was estimated by a BCA protein assay and the concentration of ATP released was expressed relative to the amount of protein.

### Statistical analysis

Data are presented as the mean ± standard error of the mean from the number of experiment (data from three or more independent experiments). Data were analyzed using Prism software (GraphPad) either by an unpaired two tailed Student’s t-test to determine difference between two groups, or one-way ANOVA with appropriate multiple-comparison test to determine difference between more than two groups.

## Supporting information

supplementary movie 1 Ollivier et al.

## Acknowledgment

This work was supported by Agence National de la Recherche (ANR-14-CEE11-0004-01) and the LabEx ICST (ANR-11-LABX-0015). We thank the iExplore animal facility (IGF, Montpellier) and the Arpege plateform (IGF, Montpellier) for the use of plate reader for cell-population fluorescence. We thank Hélène Hirbec for her assistance with statistical analysis and Etienne Audinat for reading the manuscript.

## Legends to supplementary figures

**Supplementary figure 1:**
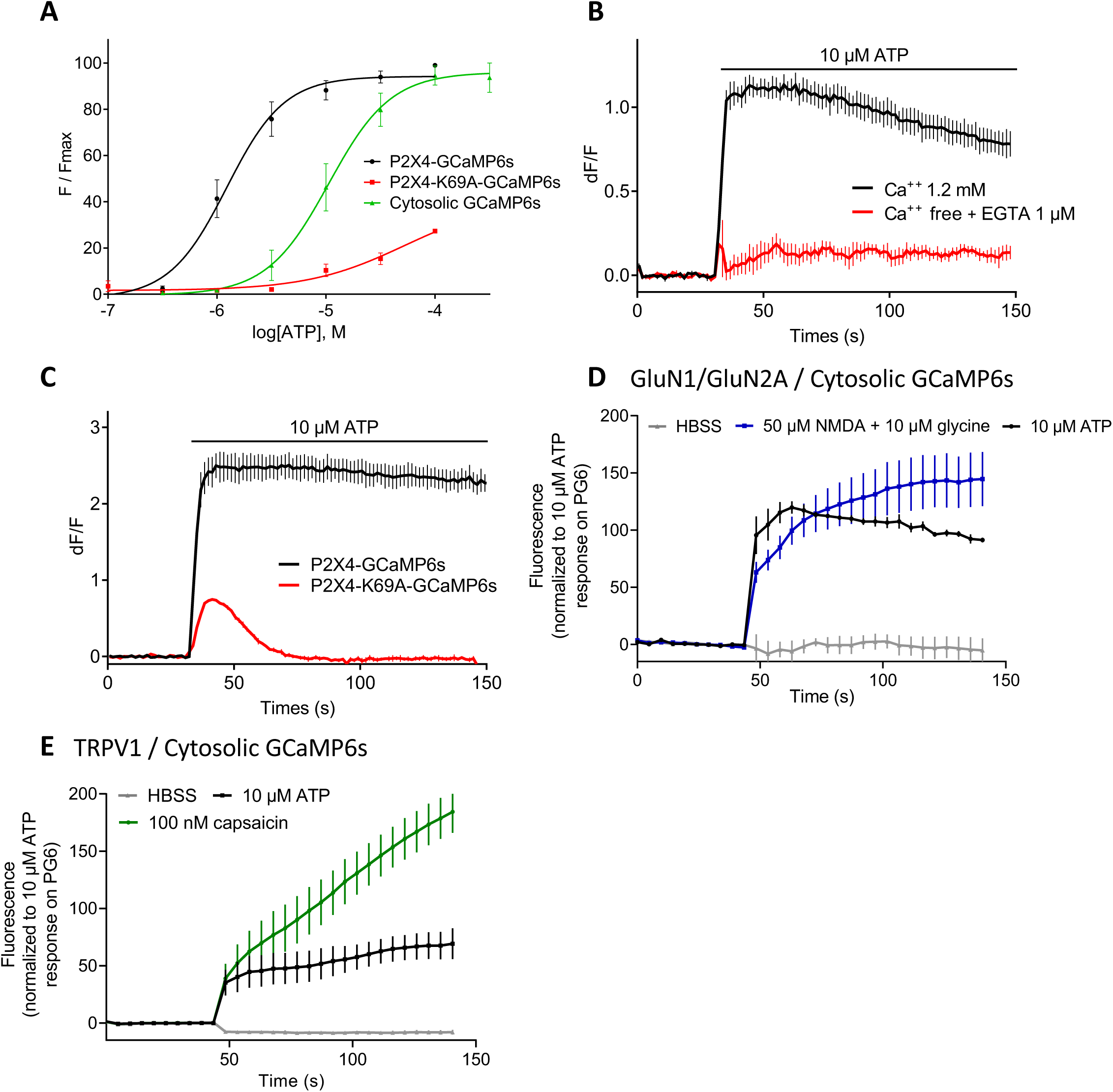
Calcium dependency of P2X-GCaMP6s. **A.** Normalized ATP dose-response curves for P2X4-GCaMP6s, P2X4-K69A-GCaMP6s and cytosolic GCaMP6s in transfected HEK cells. **B and C.** Representative traces from a plate reader of changes in dF/F values triggered by 10 μM ATP. HEK cells expressed either P2X4-GCaMP6s in the presence or the absence of extracellular calcium (B), or P2X4-GCaMP6s and P2X4-K69A-GCaMP6s (C). For all panels data are mean ± SEM of at least N > 3. **D.** Representative traces of normalized fluorescence changes evoked by ATP (10 μM, black), HBSS (gray) or NMDA/glycine (50/10 μM, blue) in HEK cells expressing NMDA receptor subunits GluN1 and GluN2A in combination with cytosolic GCaMP6s. **E.** Representative traces of normalized fluorescence changes evoked by ATP (10 μM, black), HBSS (gray) or capsaicin (100 nM, green) in HEK cells expressing TRPV1 in combination GCaMP6s. For D and E, data were generated from a plate reader with n = 3 wells per condition.

**Supplementary figure 2:**
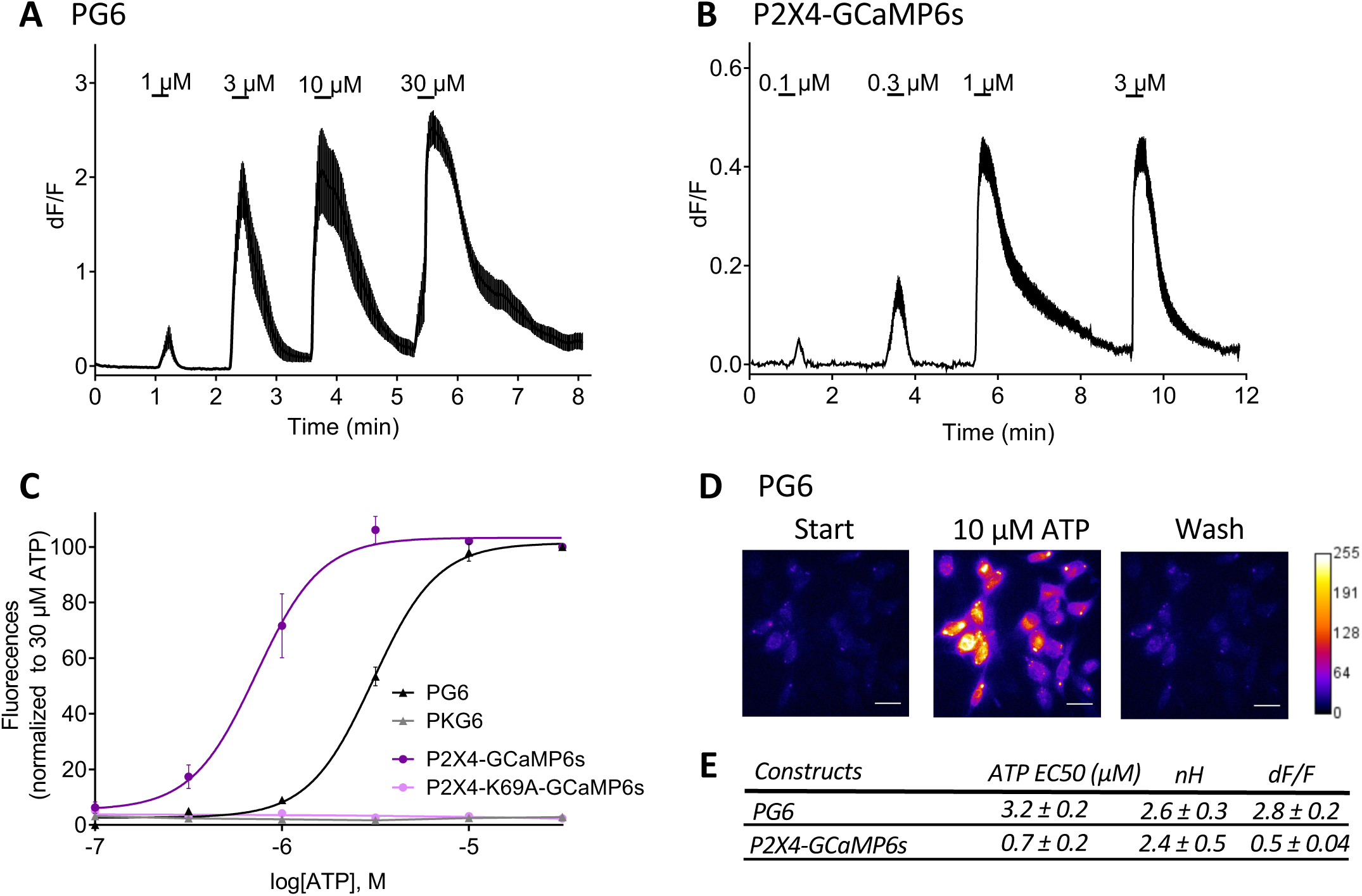
Characterization of PG6 and P2X4-GCaMP6s biosensors in the 1321N1 cell line. **A and B.** Representative traces of changes in dF/F values triggered by increasing concentration of ATP in 1321N1 cells stably expressing PG6 (A) or P2X4-GCaMP6s (B) fusions. ATP-evoked fluorescence was recorded by video microscopy (λem/exc=485/538, acquisition rate 1Hz) n = 13-17 cells **C.** Normalized concentration-response curves for ATP evoked changes in F/Fmax for PG6 or P2X4-GCaMP6s fusions and the respective mutant carrying the K69A substitution. Data are expressed as means ± SEM from n = 3 wells per condition from N > 7 independent experiments. **D.** Images showing fluorescence changes (λexc/em 485/540) of 1321N1 cells expressing PG6 before, during and after washing of ATP application (10 μM). Scale bar 20μm. **E.** Summary of EC50, Hill coefficient (nH) and maximal dF/F for PG6 and P2X4-GCaMP6s fusions expressed in 1321N1 cells.

**Supplementary figure 3:**
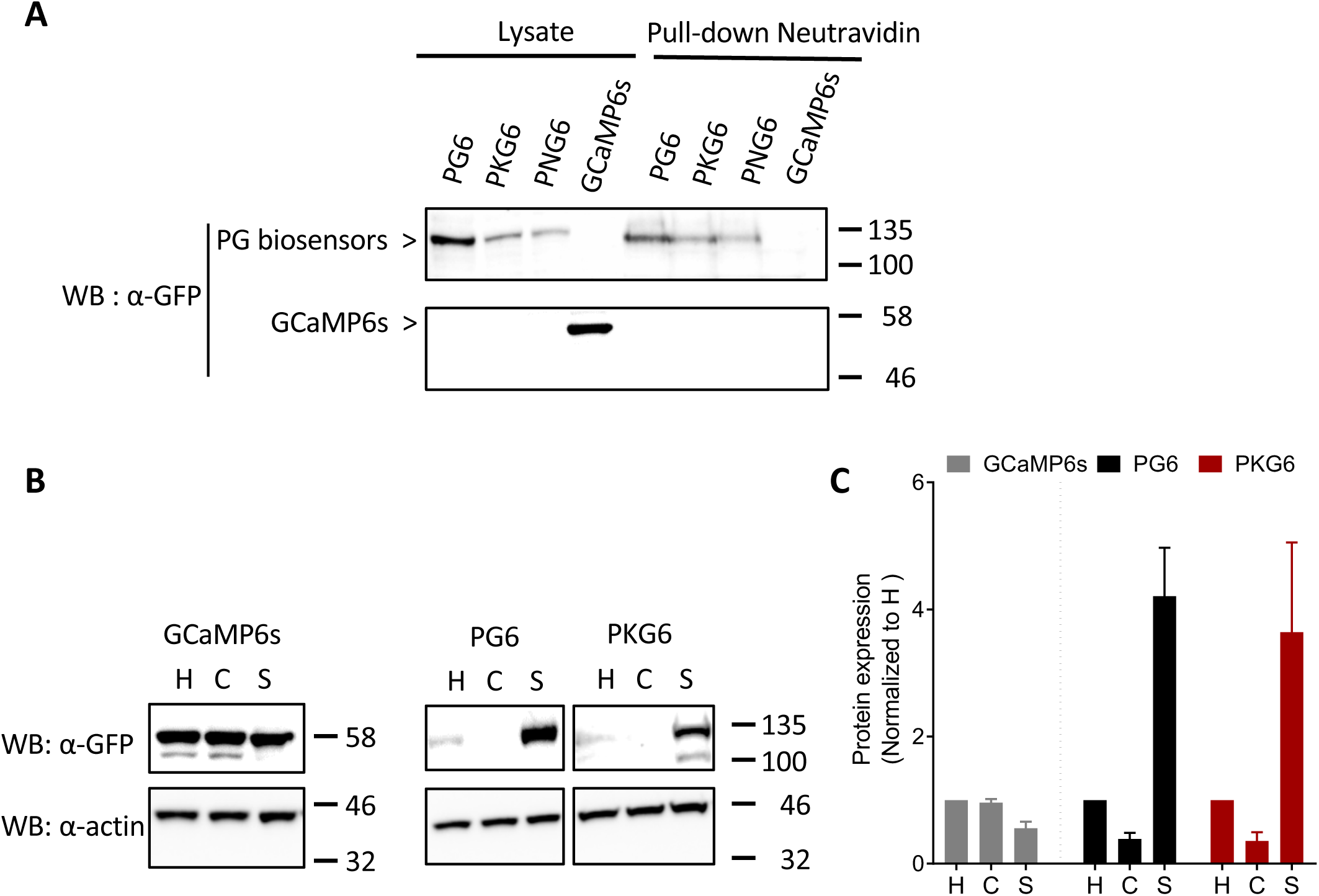
Biochemical analysis of PG6 in neurons. **A.** Analysis of cell surface expression of PG6 series by biotinylation. After biotinylation of living neurons, proteins were pulled down using neutravidin conjugated beads; biotinylated and total proteins were identified using an anti-GFP antibody. All PG6s (WT and mutants) are present at the cell surface. **B.** localization of PG6 and PKG6 in the synaptic fraction. Representative Western blot analysis of PG6, PKG6 and GCaMP6s in homogenate (H), cytosolic (C) and synaptic (S) fractions. Proteins were revealed using anti-GFP or anti-actin antibodies. PG6 and PKG6 are present in the synaptic fraction and to a lower extend in the homogenate, while cytosolic GCaMP6s is found in all fractions. **C.** Quantification of N = 6 for PG6 and PKG6, and N = 3 experiments for GCaMP6s. Results are mean ± SEM.

**Supplementary figure 4:**
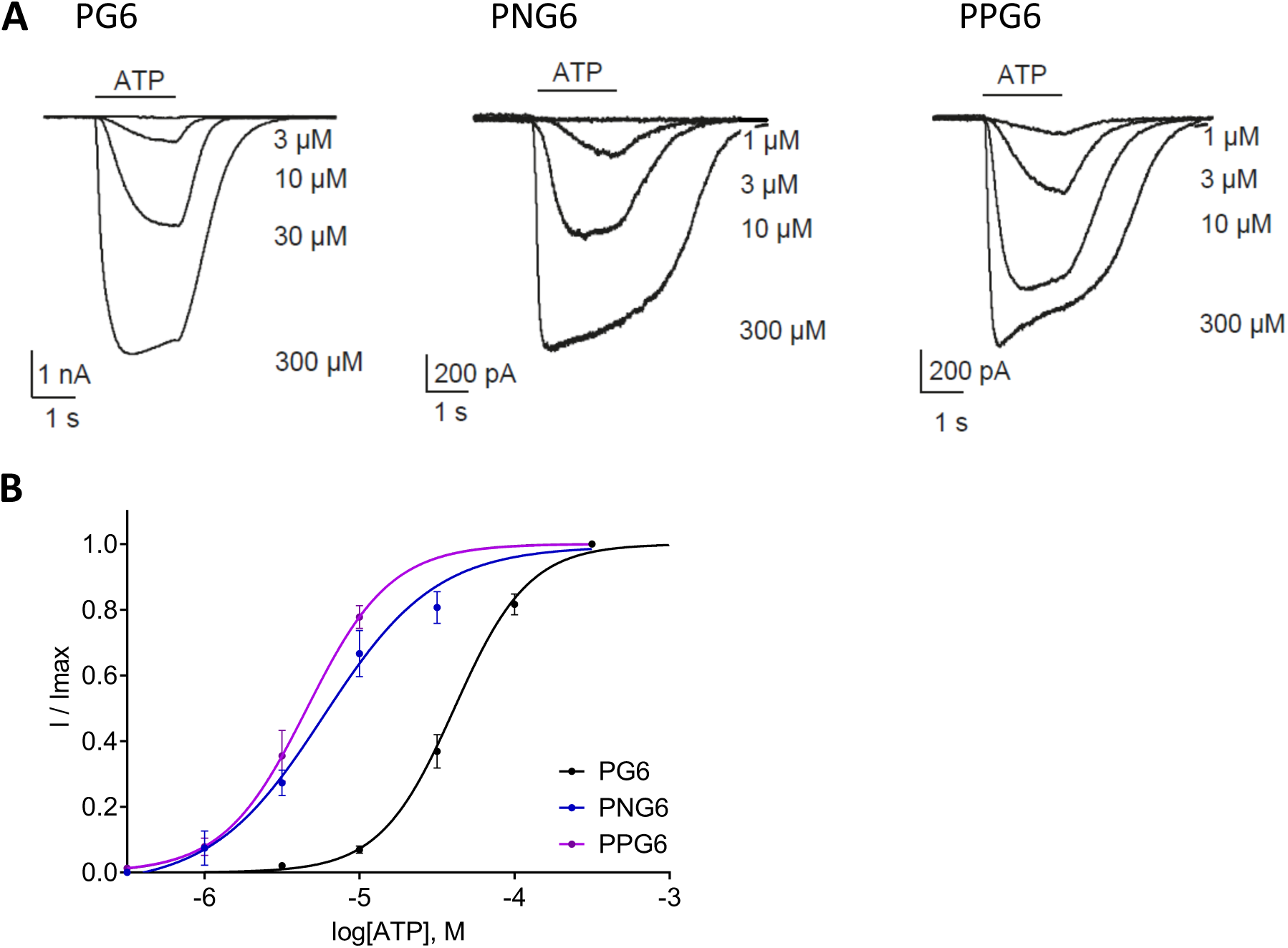
Electrophysiological characterization of PG6 high sensitivity mutants. **A and B.** Representative traces (A) and normalized dose-response curves (B) of ATP-evoked currents measured by whole-cell recording in HEK cells expressing wild type PG6, PNG6 and PPG6; data are mean ± SEM of N = 4, 3 and 4 experiments, respectively.

**Supplementary figure 5:**
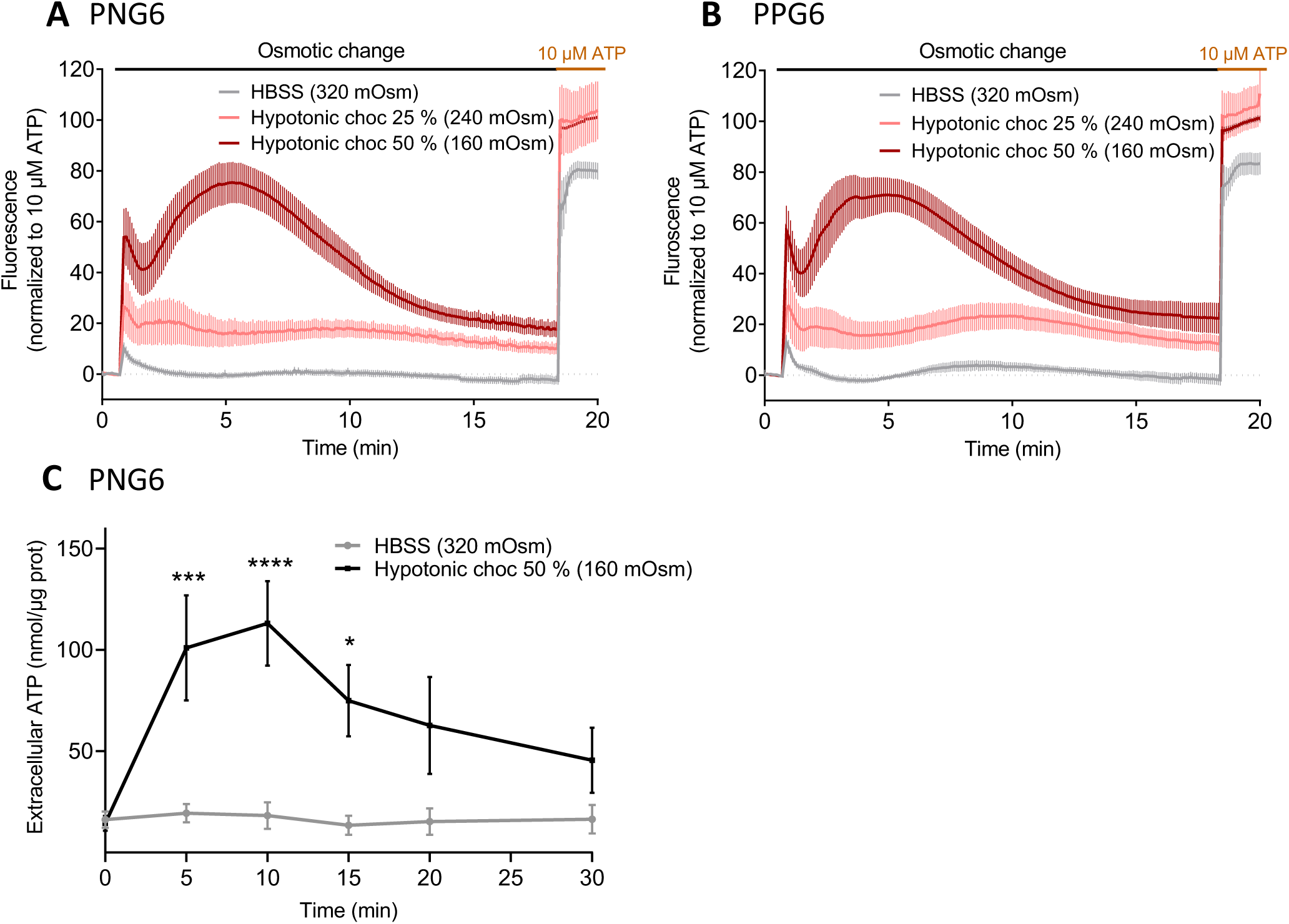
Sensing ATP release during hypotonic challenges. **A and B**. HEK cells transfected with either PNG6(A) or PPG6 (B) were challenged by medium with normal osmolarity (320 mOsm, grey) or medium with osmolarity reduced by 25 % (240 mOsm, pink) or 50 % (160 mOsm, brown). Variations of fluorescence were recorded with a plate reader. Representative results are mean ± SEM, n = 3. **C**. ATP dosage in the extracellular media of untransfected HEK cells using a bioluminescent plate reader assay (CellTiter-Glo). Extracellular ATP concentration is measured at 0, 5, 10, 15, 20, 25 and 30 min after HBBS (320 mOsm, grey) or medium with osmolarity reduced by 50 % (160 mOsm, black) application. Two-way ANOVA with Bonferroni’s multiple comparison, *, *** and **** indicate p value < 0.05, < 0.001 and < 0.0001 respectively.

**Supplementary figure 6:**
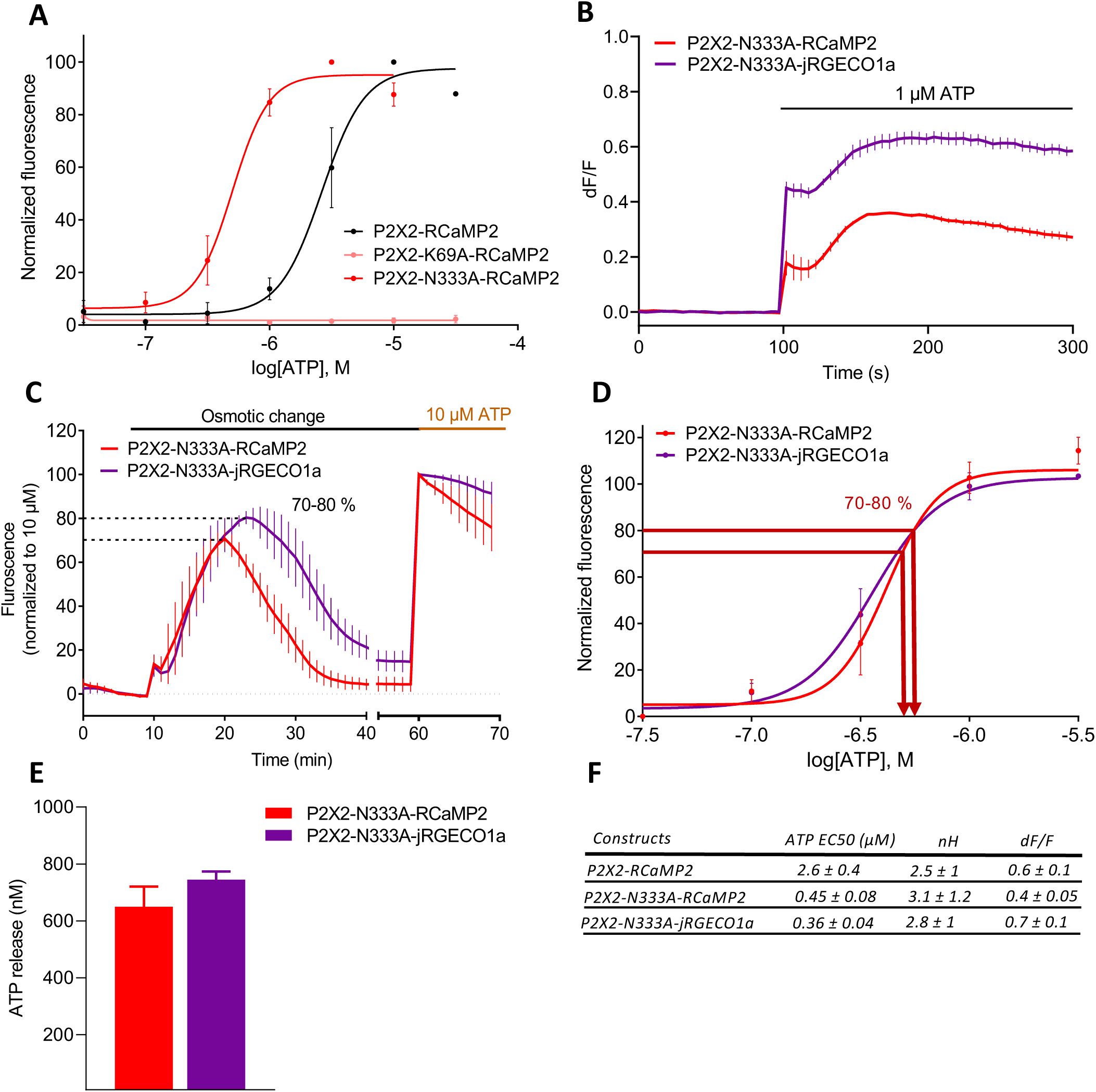
Comparison of the red-shifted ATP sensors. **A**. Pharmacological characterization of P2X2- and P2X2-N333A-RCaMP2. Normalized ATP dose responses curves were performed using a plate reader on HEK cells expressing either P2X2-, P2X2-N333A-or P2X2-K69A-RCaMP2. **B.** Representative traces of changes in dF/F values triggered after 1 μM ATP application in HEK cells expressing P2X2-N333A-RCaMP2 or P2X2-N333A-jRGECO1a. Note that the jRGECO1a-evoked fluorescence is significantly brighter than RCaMP2. Data were generated from a plate reader with n ≥ 3 wells per condition. **C.** Comparison of hypotonicity-evoked fluorescence in HEK cells expressing either P2X2-N333A-RCaMP2 or P2X2-N333A-jRGECO1a. Data were normalized to the fluorescence evoked by a 10 μM ATP application. **D.** Normalized concentration-response curves for ATP for P2X2-N333A-RCaMP2 or P2X2-N333A-jRGECO1a. Curves were generated from N = 3 independent experiments. **E.** Quantification of ATP release during a 160 mOsm hypotonic challenge of cells expressing P2X2-N333A-RCaMP2 and P2X2-N333A-jRGECO1a. N = 4 independent experiments. Data from all panels are mean ± SEM. **F**. Summary of EC50, Hill coefficient (nH) and maximal dF/F for P2X2 red shifted biosensors.

**Supplementary Movie 1: Representative movie of ATP responses in hippocampal neurons expressing PG6, related to figure 4B.** Different ATP concentrations were applied for 10 seconds at, at least, two minutes interval. Baseline fluorescence (F0) was calculated by average of the 20 initial frames. Images were pseudo-colored by F/F0 ratio and exported as JPEG and played at 31.25 fps. The timer indicate real recording time. Indications of ATP concentration are displayed during 1 minute before application (in white) and during the 10 perfusion (in yellow). Scale bar, 5 μM.

